# *picd-1*, a gene that encodes CABIN1 domain-containing protein, interacts with *pry-1/Axin* to regulate multiple processes in *Caenorhabditis elegans*

**DOI:** 10.1101/2021.09.27.462077

**Authors:** Avijit Mallick, Shane K. B. Taylor, Sakshi Mehta, Bhagwati P. Gupta

## Abstract

AXIN family members control diverse biological processes in eukaryotes. As a scaffolding protein, AXIN facilitates interactions between cellular components and provides specificity to signaling pathways. Despite its crucial roles in metazoans and discovery of a large number of family members, the mechanism of AXIN function is not very well understood. The *C. elegans* AXIN homolog PRY-1 provides a powerful tool to identify interacting genes and downstream effectors that function in a conserved manner to regulate AXIN-mediated signaling. Previous work demonstrated *pry-1*’s essential role in developmental processes such as reproductive system, seam cells, and a P lineage cell P11.p. More recently, our lab carried out a transcriptome profiling of *pry-1* mutant and uncovered the essential role of the gene in lipid metabolism, stress response, and aging. In this study, we have extended the work on *pry-1* by reporting a novel interacting gene *picd-1* (***p****ry-1*-**i**nteracting **C**ABIN1 **d**omain containing). Our findings have revealed that *picd-1* plays an essential role in *C. elegans* and is involved in several *pry-1*-mediated processes including regulation of stress response and lifespan maintenance. In support of this, *picd-1* expression overlaps with *pry-1* in multiple tissues throughout the lifespan of animals. Further experiments showed that *picd-1* inhibits CREB-regulated transcriptional coactivator homolog CRTC-1 function, which promotes longevity in a calcineurin-dependent manner. These data provide evidence for an essential role of the CABIN1 domain protein PICD-1 in mediating PRY-1 signaling in *C. elegans*.

## INTRODUCTION

Signaling pathways confer the ability on cells to communicate with each other and with their environment. Because of their essential roles, activities of pathway components are regulated via interactions with a host of cellular factors. Scaffolding proteins such as Axin family are a group of proteins that bring together different proteins to facilitate interactions, which regulate their activity ^1^. Axin was initially discovered as a negative regulator of WNT-mediated signaling cascade, but subsequent work revealed a much broader role of family members in other pathways ^1–3^.

In the nematode *C. elegans*, the Axin homolog PRY-1 controls processes such as embryogenesis, neuronal differentiation, vulval development, P11.p cell fate, and seam cell development ^1,4–6^. More recently, work from our lab has shown that PRY-1 is also essential for lipid metabolism, stress response, and lifespan maintenance^7–9^. While WNT-dependent function of PRY-1, e.g., in vulval cells, involves its interactions with APR-1 (APC family) and GSK-3 (GSK3β), leading to phosphorylation of BAR-1 (β-Catenin) ^4^, little is understood about factors that interact with PRY-1 in non-WNT-dependent processes.

A thorough understanding of PRY-1 function not only requires identification of interacting proteins but also its downstream effectors. We earlier reported a transcriptome profiling of PRY-1, which revealed many differentially expressed genes involved in lipid regulation and aging ^7^. In this study, we report characterization of a novel downstream effector of *pry-1* signaling, namely *picd-1* that plays essential roles in regulating multiple developmental and post-developmental processes. PICD-1 shares a domain with the mammalian calcineurin-binding protein 1 (CABIN1). CABIN1 negatively regulates calcineurin signaling, the pathway known to affect a wide array of cellular functions including stress response and lifespan ^10–13^. Our data suggests that PICD-1 negatively regulates CREB-regulated transcriptional coactivator (CRTC) homolog, CRTC-1, function to promote longevity mediated by calcineurin signaling ^14^. These results form the basis of future investigations to understand the mechanism of PICD-1 function in PRY-1-mediated signaling.

## RESULTS

### *picd-1* encodes a CABIN1 domain containing protein

During a CRISPR-based screen to isolate alleles of *pry-1*, we recovered a secondary mutation (*gk3701*) in F56E10.1, now named as *picd-1* (***p****ry-1* **i**nteracting **C**ABIN1 **d**omain containing, see Methods). The *pry-1(gk3681); picd-1(gk3701)* double mutants exhibit a significant increase in Pvl phenotype (77%, compared to 66% in *pry-1* mutants alone) and pronounced protrusions that result into frequent bursting at the vulva (**Table 1, Figure 1A and B)**. Sequence analysis of PICD-1 identified orthologs in other nematode species (**Figure 1C**), all of which contain a domain similar to Histone transcription regulator 3 (Hir3)/Calcineurin-binding protein (CABIN1) family members (IPR033053, https://www.ebi.ac.uk/interpro/) (**Figure 1C and D**). Alignments of PICD-1 with the human CABIN1 (isoform a) showed 26% (729/2853) identity and 38% (1080/2853) similarity (EMBOSS stretcher pairwise alignment tool, https://www.ebi.ac.uk/Tools/psa/). Human CABIN1 is known to be part of a histone H3.3 chaperone complex, HUCA (HIRA/UBN1/CABIN1/ASF1a) that is involved in nucleosome assembly. Likewise, Gene Ontology analysis (www.wormbase.org) identified that *picd-1* is associated with the biological process ‘DNA replication-independent nucleosome assembly’ (GO:0006336) and the cellular component ‘nucleus’ (GO:0005634). Thus, *picd-1* is likely to encode a nuclear protein with function in chromatin assembly and regulation of gene expression. In support of this, in silico analysis revealed that PICD-1 contains 49 amino acid residues predicted to bind DNA (http://biomine.cs.vcu.edu/servers/DRNApred/#References ^15^) (**Table S1**).

**Table 1:** Analysis of VPC induction, P12.pa cell fate, Pvl, and Muv penetrance in different strains. *Extra P12.pa cell in the place of P11.p.

| Genotype | VPC induction pattern (% induced) |  |  |  |  |  |  | Over-induced vulva and P12.pa |  |  | Pvl and Muv |  |  |
| --- | --- | --- | --- | --- | --- | --- | --- | --- | --- | --- | --- | --- | --- |
|  | P3p | P4p | P5p | P6p | P7p | P8p | N | % over-induced | % P12pa* | N | % Pvl | %Muv | N |
| N2 | 0 | 0 | 100 | 100 | 100 | 0 | 20 | 0 | 0 | 20 | 0 | 0 | 20 |
| *pry-1(gk3682)* | 27.3 | 4.5 | 100 | 100 | 86.4 | 4.5 | 22 | 36.4 | 72.7 | 22 | 65.7 | 31.3 | 40 |
| *pry-1(mu38)* | 18.2 | 0 | 100 | 100 | 81.8 | 9.1 | 22 | 27.2 | 81.8 | 22 | 59.6 | 34.2 | 50 |
| *picd-1(gk3701)* | 0 | 0 | 100 | 100 | 100 | 0 | 20 | 0 | 0 | 20 | 5 | 0 | 40 |
| *picd-1(bh40)* | 0 | 0 | 100 | 100 | 100 | 0 | 20 | 0 | 0 | 20 | 16.7 | 0 | 40 |
| *pry-1(gk3681);*<br>*picd-1(gk3701)* | 21.7 | 8.7 | 100 | 100 | 87 | 13 | 23 | 39.1 | 74 | 23 | 76.8 | 22 | 40 |
| *pry-1(mu38);*<br>*picd-1(bh40)* | 12.5 | 12.5 | 100 | 100 | 92 | 12.5 | 24 | 25 | 100 | 24 | 80 | 20 | 40 |

**Figure 1:**
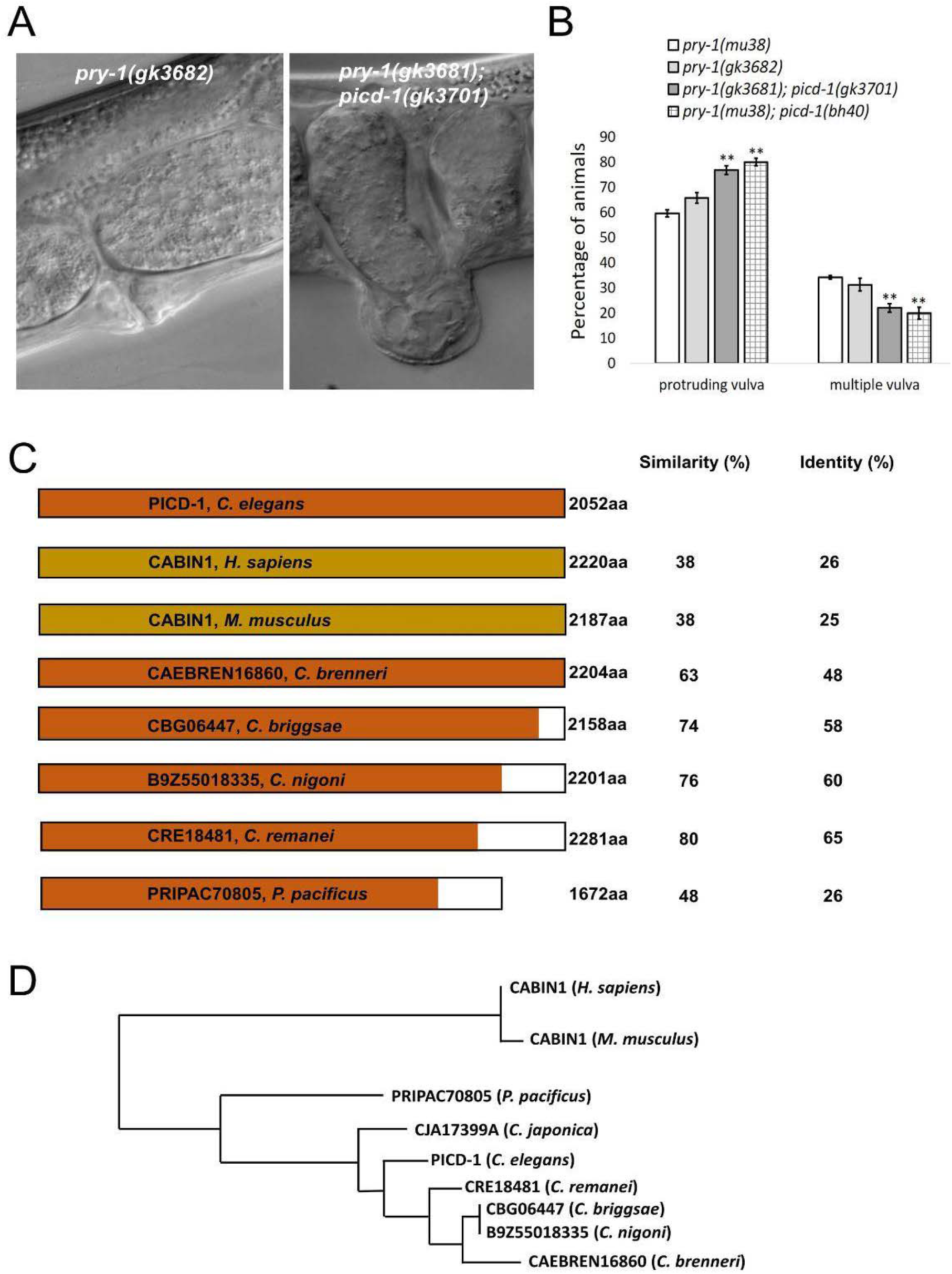
PICD-1 has sequence similarity with mammalian CABIN1 protein. **(A)** Representative images of Pvl phenotype in *pry-1(gk3682)* and *pry-1(gk3681); picd-1(gk3701)* animals. **(B)** *picd-1* mutation enhances Pvl phenotype of *pry-1* mutants. Data represent a cumulative of two replicates (n > 30 animals) and error bars represent the standard deviation. Statistical analyses for panel (**B)** were done using one-way ANOVA with Dunnett’s post hoc test and significant differences are indicated by stars (*): ** (*p* <0.01). **(C)** Protein sequence comparison of PICD-1 with mammalian CABIN1 and homologs in the nematodes. **(D)** Multiple sequence alignment dendrogram generated by LIRMM (http://www.phylogeny.fr/simple_phylogeny.cgi) using default parameters.

### *picd-1* is expressed in multiple tissues

To gain further insights into the function of *picd-1*, we created transgenic animals carrying *picd-1p::GFP* transcriptional reporter. The analysis of transgenic animals revealed GFP fluorescence during development in tissues such as pharynx, intestine, body wall muscles, hypodermis (seam cells), gonad, and vulva **(Figure 2)**. This pattern of expression resembles that of *pry-1* recently described by our group ^8^. As *picd-1::GFP* animals entered adulthood, fluorescence was localised to the intestine and certain head neurons **(Figure 2)** which persisted throughout the life of animals (data not shown). A broad range of *picd-1* expression is also supported by previously published RNA-seq and microarray studies^16,17^. Overall, our expression analysis suggests that *picd-1* functions in multiple tissues and may play roles in *pry-1*-mediated processes. Data presented in the following sections support this conclusion.

**Figure 2:**
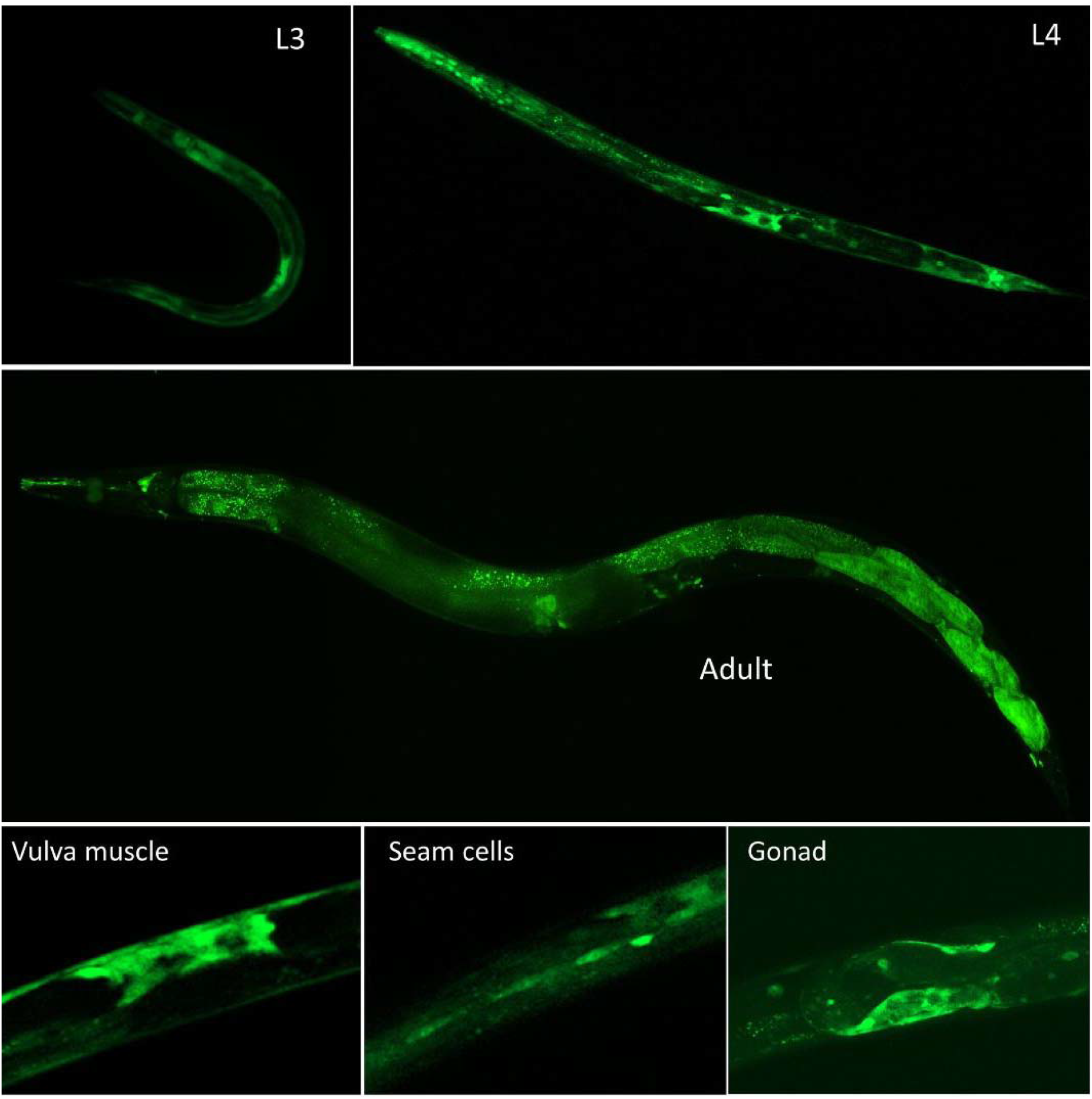
Reporter analysis of *picd-1* reveals expression in multiple tissues. Representative images of animals expressing *picd-1* in both larval and adult stages. Tissues that express *picd-1* include pharynx, gonad, head neurons, intestine, vulva and body wall muscle.

### Mutations in *picd-1* cause Pvl and Egl defects

In addition to using the *gk3701* strain to examine mutant phenotypes, we generated a new allele *bh40* that contains multiple in-frame stop codons in exon 1 (**see Methods and Figure 3A and B**). qPCR analysis showed that *bh40* and *gk3701* greatly reduce *picd-1* transcript levels (**Figure 3C**). Interestingly, while the Pvl phenotype of *pry-1(mu38)* is enhanced by both alleles (**Figure 1A and B**), neither has an obvious impact on the Muv penetrance of *pry-1*(mu38) animals. In fact, the double mutants are slightly less Muv compared to the *pry-1(mu38)* alone (**Table 1, Figure 1B**), which may be due to morphogenetic defects since vulval induction is not affected by any of the *picd-1* mutations (**Figure 3D, Table 1**). Similar phenotypes are also observed following *picd-1* RNAi (**Figure S1**).

**Figure 3:**
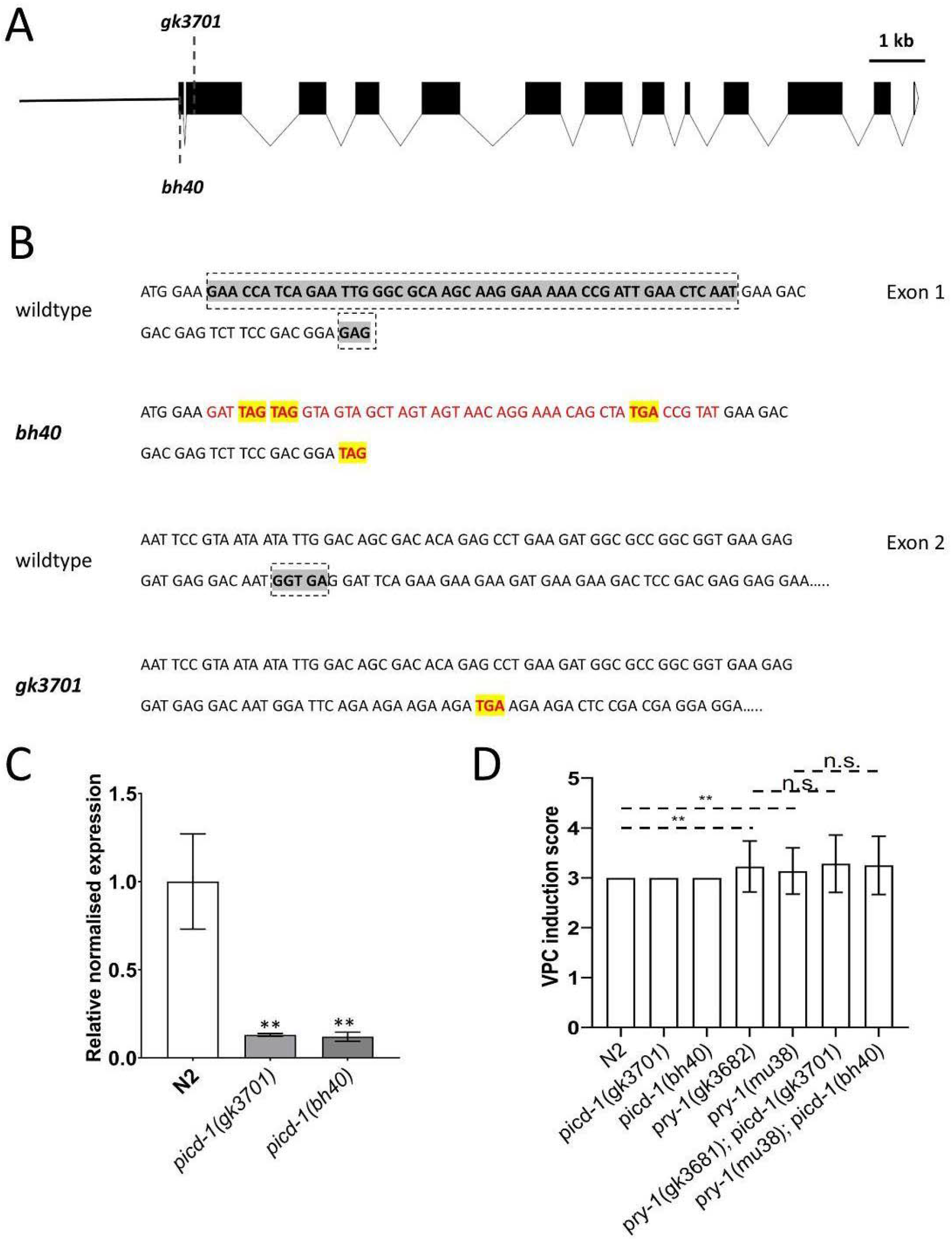
Mutation in *picd-1* enhances *pry-1* Pvl defect. **(A)** Schematic diagram of the *picd-1* coding transcript showing the location of two different alleles: *bh40* (first exon) and *gk3701* (second exon). **(B)** Highlighted sequences show regions of the wild-type sequence (grey color) that have been either substituted (red color) or deleted in the respective mutant alleles. Newly introduced in-frame stop codons are marked in yellow. **(C)** Bar graph showing VPC induction score in *picd-1* and *pry-1* mutants alone, and *pry-1; picd-1* double mutants compared to N2. Data represent the mean of two replicates and the error bar showing the standard deviation (n > 15 animals per batch). Statistical analyses was done using one-way ANOVA with Dunnett’s post hoc test and significant differences are indicated by stars (*): ** (*p* <0.01).**(D)** Expression level of *picd-1* in *pry-1(gk3682)* and *pry-1(mu38)* mutants at the L1 stage compared to wild-type. Data represent the mean of two replicates and error bars represent the standard error of mean. Significance was calculated using Bio-Rad software (one-way ANOVA) and significant differences are indicated by stars (*): ** (*p* <0.01).

Phenotypic analysis of both *picd-1* mutant strains revealed that animals do not show any obvious sign of sickness. Upon careful plate-level examination, we found that the gene is involved in the development of the egg laying system. While both alleles show weak Pvl phenotypes on their own, i.e., independent of *pry-1* mutations, *bh40* appears to be a stronger loss of function allele (*gk3701*: 5% Pvl and *bh40*: 16.7%) (**Table 1**). Furthermore, *picd-1(bh40),* but not *picd-1(gk3701),* animals exhibit abnormal vulval invagination (**Figure 4A**), indicating that defects in morphogenetic processes may contribute to enhanced Pvl phenotype of mutant animals. We also observed that the vulval morphology phenotype of *bh40*, but not *gk3701*, was dominant over *pry-1(mu38)* **(Figure 4A)**. Among other defects, we observed that *picd-1(gk3701)* worms lay eggs normally, but *picd-1(bh40)* are weakly Egl (**Figure S2, Video 1**) and both Pvl and Egl phenotypes of *picd-1(bh40)* animals are enhanced when grown at 25°C (**Figures 4B and S2**).

**Figure 4:**
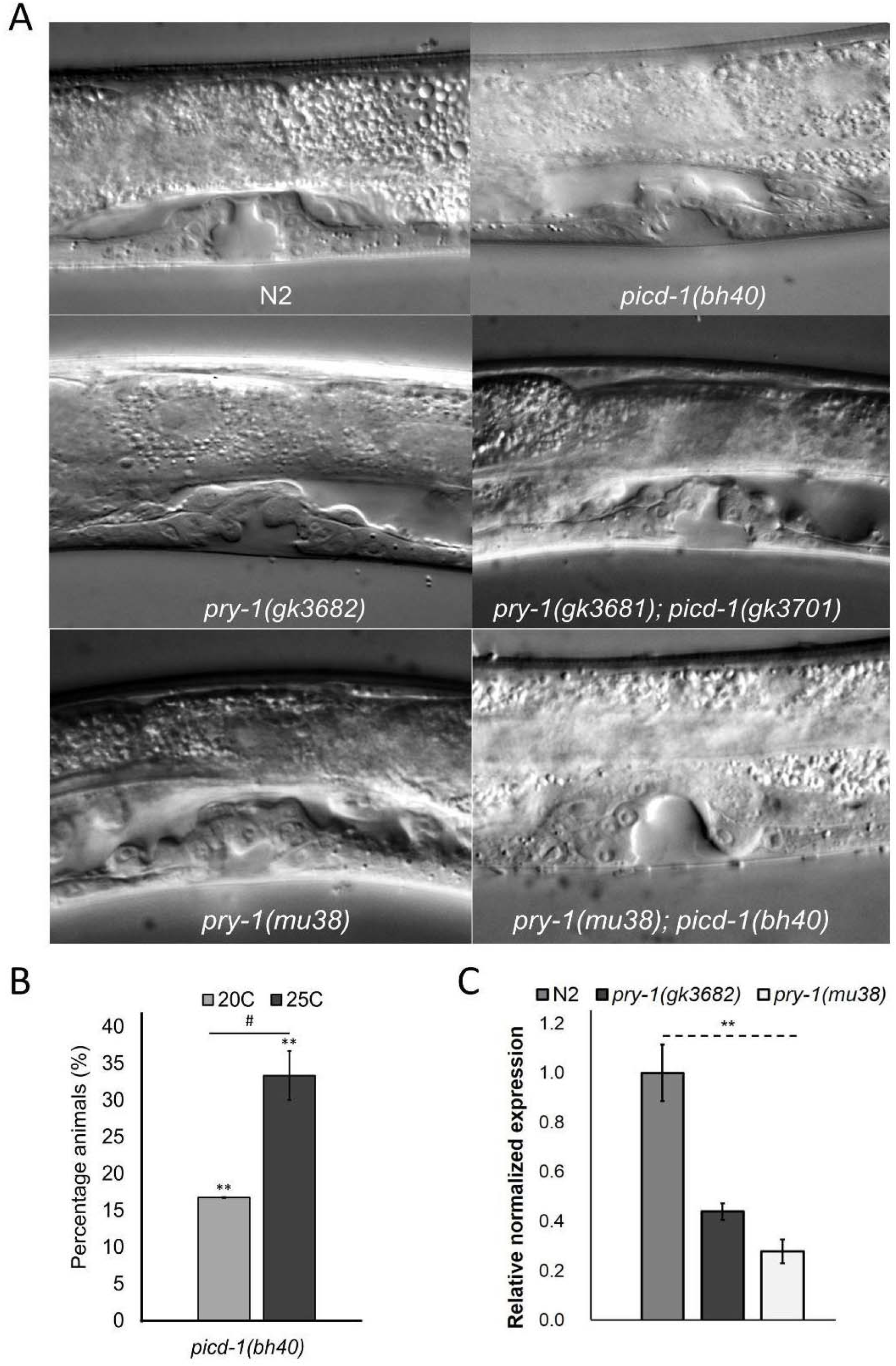
*picd-1* regulate vulva morphology. **(A)** Representative vulva morphology images of wild-type, *picd-1(bh40), pry-1(gk3682), pry-1(gk3681); picd-1(gk3701), pry-1(mu38)* and *pry-1(mu38); picd-1(bh40)* animals at mid-L4 stage. **(B)** Bar graph showing the percentage of *picd-1(bh40)* mutants showing Pvl phenotype at 20C and 25C compared to wild-type animals (which show no Pvl) (Also see Table 1). **(C)** Expression level of *picd-1* in both the *pry-1* mutants. Data represent the mean of two replicates and error bars represent the standard error of mean. Significance was calculated using Bio-Rad software (one-way ANOVA) and significant differences are indicated by stars (*): ** (*p* <0.01). Data represent a cumulative of two replicates (n > 30 animals) and error bars represent the standard deviation. Statistical analyses was done using one-way ANOVA with Dunnett’s post hoc test and significant differences are indicated by stars (*): ** (*p* <0.01).

Since *picd-1::GFP* pattern overlaps with that of *pry-1* and *picd-1* mutation enhances *pry-1* Pvl phenotype, we examined whether *pry-1* affects *picd-1* expression. qPCR experiments showed that *picd-1* level is drastically reduced in *pry-1* mutants (**Figure 4C**). Overall, results described in this section lead us to conclude that *picd-1* is required for the development of the reproductive system and functions genetically downstream of *pry-1*.

### *picd-1* mutations worsen the phenotypes of *pry-1* mutants

We also investigated the involvement of *picd-1* in other *pry-1*-mediated developmental and post-developmental processes. *picd-1* mutants are slow growing and take longer time to reach adulthood compared to either wildtype or *pry-1(mu38)* animals (**Figure 5A**). It was found that the slower growth phenotype of *pry-1* and *picd-1* double mutants is significantly worse than either single mutant (**Figure 5A**).

**Figure 5:**
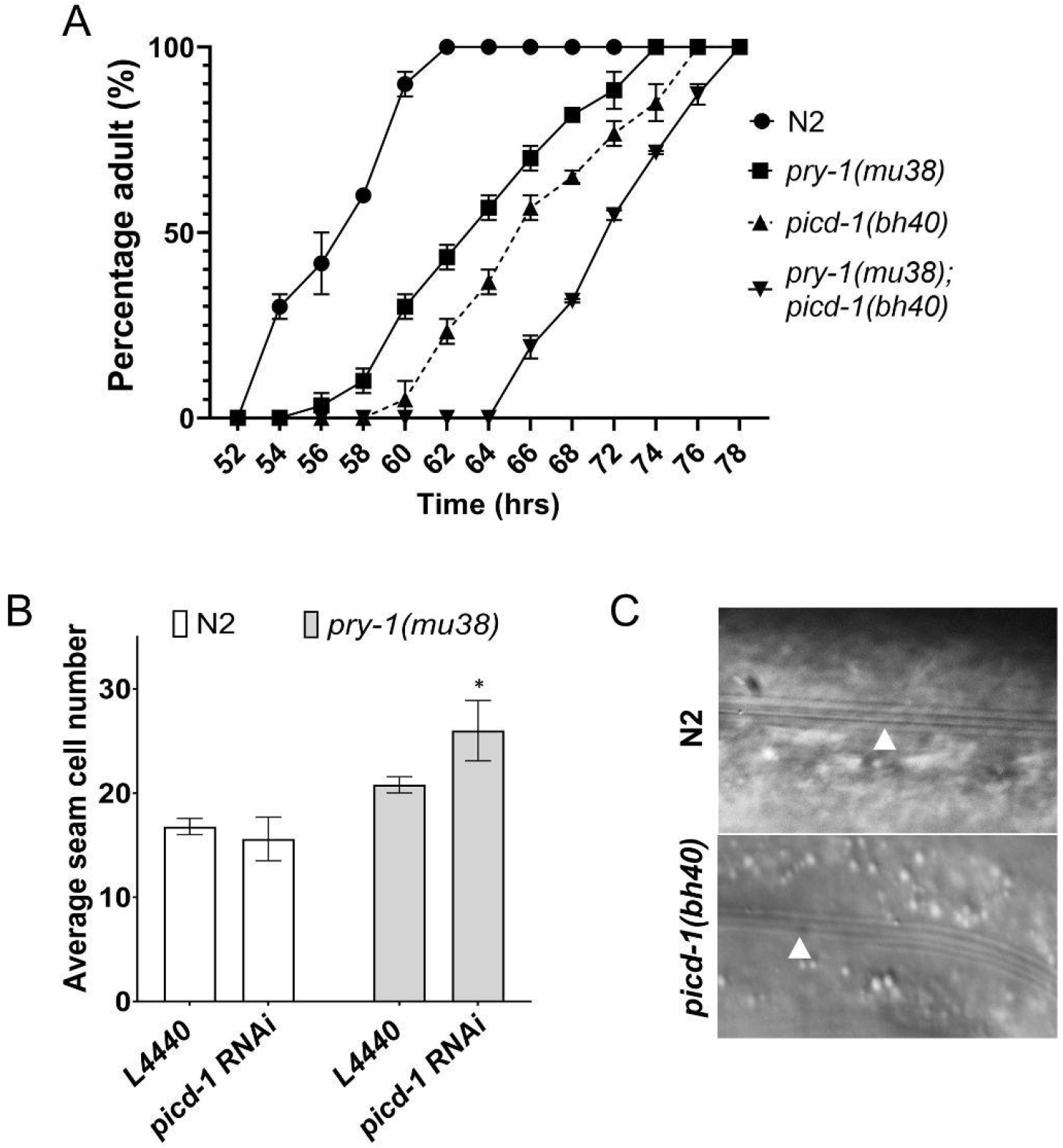
*picd-1* interacts with *pry-1* to regulate developmental timing, seam cell division and alae formation. (A) *picd-1* mutants exacerbate the developmental delay of *pry-1* mutants. Data shows the average (two replicates, n > 60 animals) time taken by *picd-1(bh40), pry-1(mu38)* and *pry-1(mu38); picd-1(bh40)* double mutants to reach adulthood compared to wild-type animals. **(B)** Bar graph showing average number of seam cells in the wild-type and *pry-1(mu38)* animals following control (L4440) and *picd-1* RNAi. Data represent a cumulative of two replicates (n > 30 animals) and error bars represent the standard deviation. Statistical analyses were done using one-way ANOVA with Dunnett’s post hoc test and significant differences are indicated by stars (*): * (*p* <0.05). **(C)** Representative images showing abnormal alae (alae shown with white arrow; extra alae shown with *) in *picd-1(bh40)* mutants compared to wild-type animals.

Mutations in *picd-1* enhance developmental defects of *pry-1(mu38)* animals as well that include a P lineage cell P11.p and seam cells. While 70-80% of *pry-1* mutants showed an extra P12.pa cell in the place of P11.p, the phenotype was fully penetrant in *picd-1(bh40)*; *pry-1(mu38)* double **(Table 1)**. The seam cell defect in *pry-1* mutants is caused by changes in asymmetric cell divisions at the L2 stage (Gleason et al 2010, Mallick et al 2019). While RNAi knockdown of *picd-1* showed no obvious seam cell defect, it enhanced the phenotype of *pry-1(mu38)* animals (**Figure 5B**). Moreover, both *picd-1* and *pry-1* mutants exhibited defects in alae, structures that are formed by differentiated seam cells (**Figure 5C**) (Mallick et al 2019). These data show that *picd-1* interacts with *pry-1* to affect P11.p and seam cell development.

In addition to developmental defects, we observed several other post-developmental abnormalities in *picd-1* mutant animals. The analysis of brood size revealed defects in *picd-1(bh40)* but not in *picd-1(gk3701)* animals (**Figure 6A and B**). While *bh40* allele does not affect embryonic viability, it drastically enhances the low brood count and embryonic lethality of *pry-1* mutants (*p* < 0.001) (**Figure 6A-C**). We also analyzed gonad morphology and oocytes and found that neither allele affects these phenotypes. Interestingly *pry-1(gk3681); picd-1(gk3701)* and *pry-1(mu38)*; *picd-1(bh40)* double mutants showed abnormal oocytes and gonads; respectively (**Figure 7A-C**). More specifically, 46 +/− 6% (n=45) of *pry-1(mu38)*; *picd-1(bh40)* animals lacked oocytes in the posterior gonad arm (**Figure 7C and D**). No such phenotype was observed in either single mutant.

**Figure 6:**
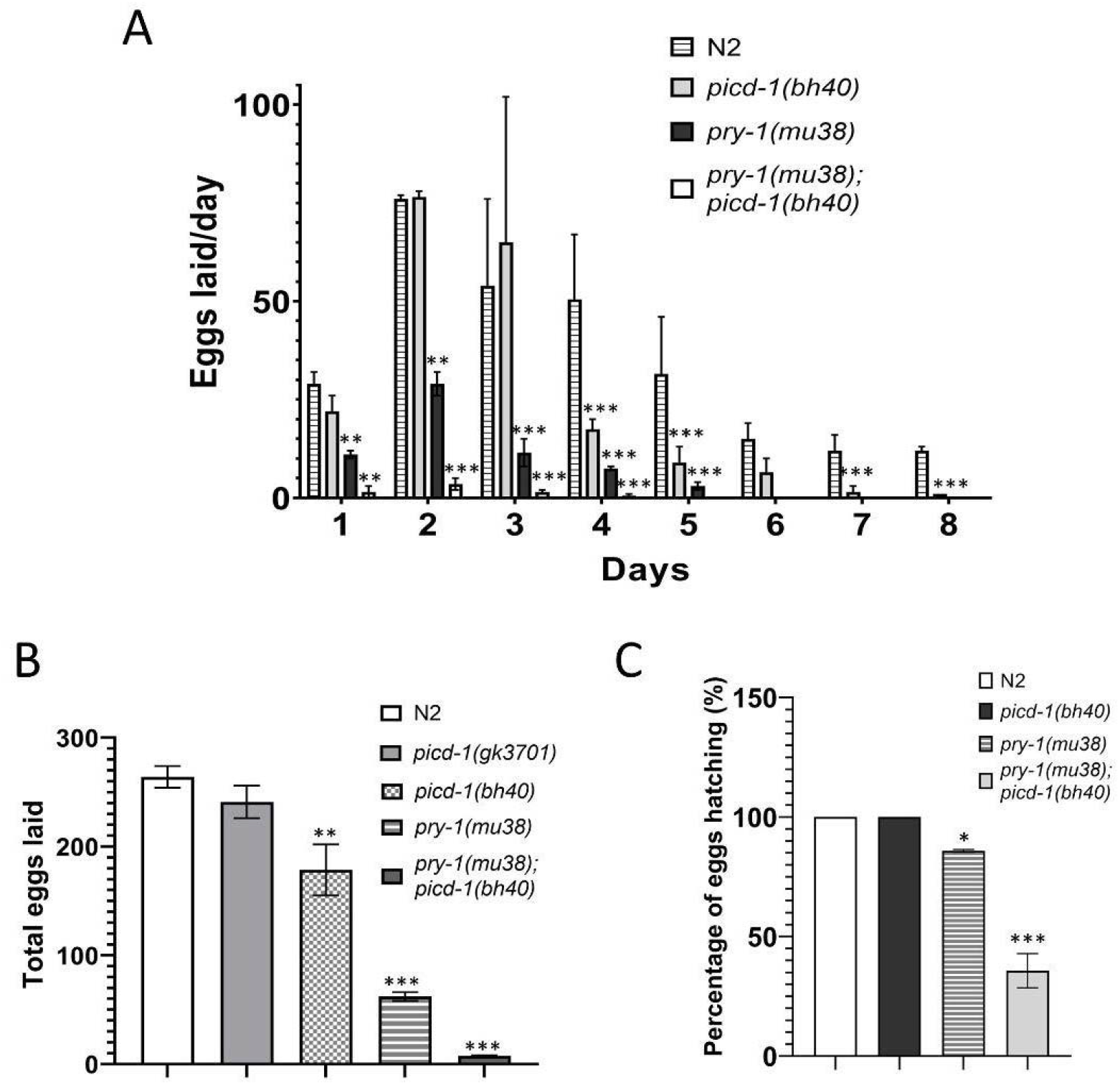
*picd-1* interact with *pry-1* to regulate brood size and embryonic survivability. **(A)** Bar graphs showing eggs laid on each day by *picd-1(bh40)*, *pry-1(mu38)* and *pry-1(mu38); picd-1(bh40)* double mutants compared to control animals over a 5 day period. **(B)** Bar graphs showing the totals number of eggs laid by *picd-1(gk3701)*, *picd-1(bh40)*, *pry-1(mu38)* and *pry-1(mu38); picd-1(bh40)* double mutants compared to control animals. **C)** Bar graph showing the percentage of the total eggs hatching for *picd-1(bh40)*, *pry-1(mu38)* and *pry-1(mu38); picd-1(bh40)* double mutants compared to control animals. Data represent a cumulative of two replicates (n > 30 animals) and error bars represent the standard deviation. Statistical analyses was done using one-way ANOVA with Dunnett’s post hoc test and significant differences are indicated by stars (*): * (*p* <0.05), ** (*p* <0.01), *** (*p* <0.001).

**Figure 7:**
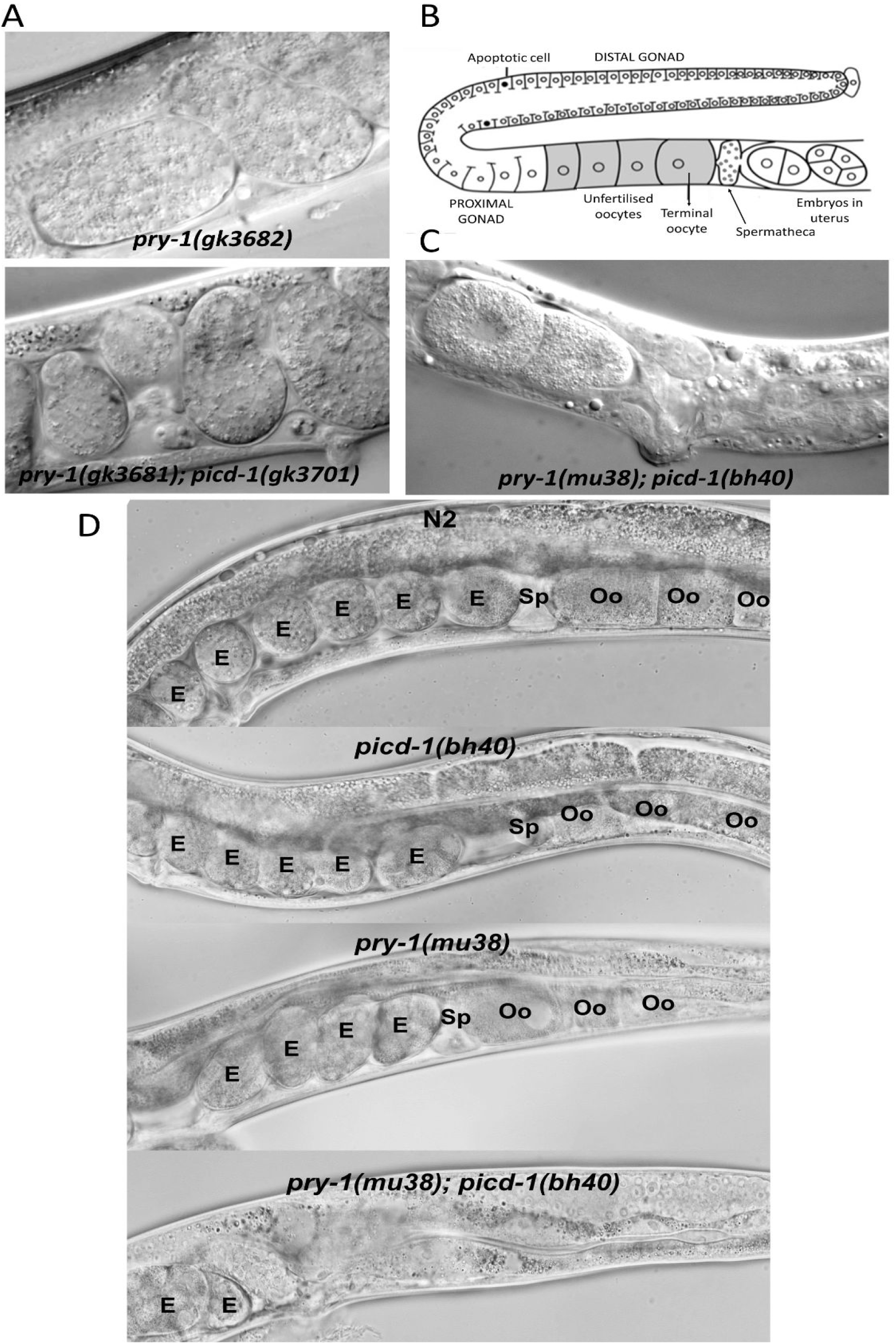
*picd-1* interacts with *pry-1* to regulate oocyte development. **(A)** *pry-1; picd-1* double mutants show abnormal oocytes and embryos morphology. **(B)** Schematic summary of major changes in the anatomy of the hermaphrodite gonad (Guardia et al 2016). **C)** Representative image of the *pry-1(mu38); picd-1(bh40)* animals with no oocyte formation in the proximal gonad arm (phenotype penetrance of 44%, also see Supplementary Figure). **(D)** Representative images of wildtype, *picd-1(bh40), pry-1(mu38)* and *pry-1(mu38); picd-1(bh40)* animals at adult stage. The spermatheca (Sp), embryos (E), and oocytes (Oo) are marked.

### *picd-1* mutants are sensitive to stress and exhibit a short lifespan

We reported earlier that *pry-1* plays a role in stress response maintenance ^6,8^. It was found that all three heat shock chaperons, i.e., *hsp-4* (ER-UPR), *hsp-6* (MT-UPR), and *hsp-16.2* (cytosolic heat shock response, HSR) are upregulated in *pry-1* mutant animals (**Figure 8A**). *picd-1* mutants showed increased expression of two of these, *hsp-4* and *hsp-16.2*, and the oxidative stress response gene *sod-3* (**Figure 8B**). Consistent with these results, both *pry-1* and *picd-1* mutants show electrotaxis defects (**Figure 8C**), a phenotype observed in animals due to abnormalities in UPR signaling^18^.

**Figure 8:**
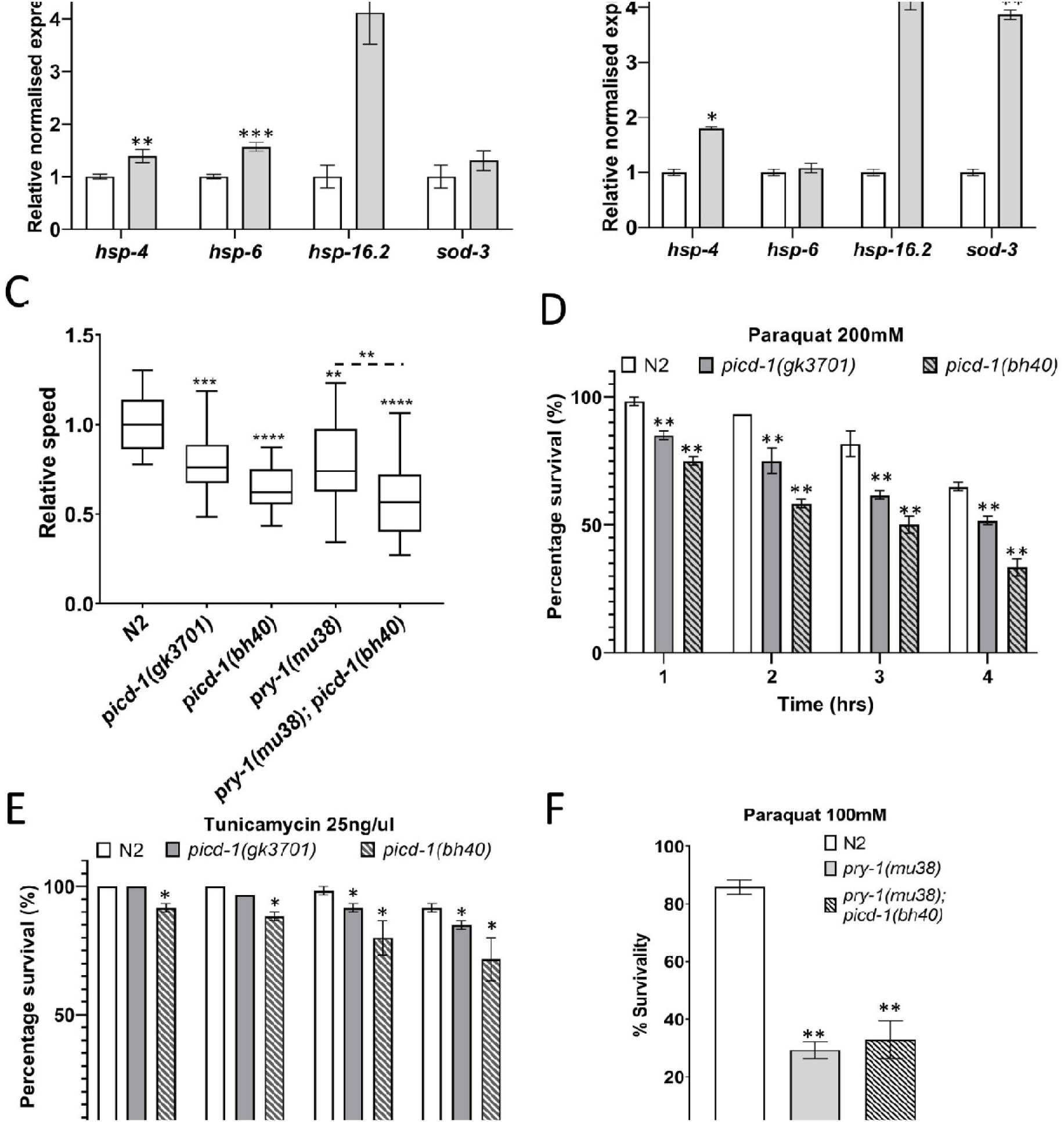
*picd-1* mutants are stress sensitive. **(A-B)** Expression levels of *hsp-4, hsp-6, hsp-16.2*, *sod-3* and *hsf-1* in *picd-1(bh40)* and *pry-1(mu38)* young adult animals. Data represent the mean of two replicates and error bars represent the standard error of mean. Significance was calculated using Bio-Rad software (one-way ANOVA) and significant differences are indicated by stars (*): ** (*p* <0.01). **(C)** Box and whisker plots represent average speed of *picd-1(gk3701)*, *picd-1(bh40)*, *pry-1(mu38)* and *pry-1(mu38); picd-1(bh40)* mutants compared to wild-type. **(D)** Bar graphs represent percentage survival of animals following paraquat exposure of 200mM over 4 hrs. (**E)** Bar graphs represent percentage survival of animals following tunicamycin exposure of 25ng/μl over 4 hrs. **(F)** Bar graphs represent percentage survival of animals following paraquat exposure of 100mM after 1 hr. For panels **C-F**, data represent a cumulative of two replicates (n > 30 animals) and error bars represent the standard deviation. Statistical analyses were done using one-way ANOVA with Dunnett’s post hoc test and significant differences are indicated by stars (*): * (*p* <0.05), ** (*p* <0.01).

To further investigate the stress sensitivity of animals lacking *picd-1* function, we examined survivability following chemical treatments. Both *gk3701* and *bh40* mutants were sensitive to paraquat and tunicamycin although the effect was more pronounced following paraquat exposure **(Figure 8D and E)**. Interestingly, *bh40* did not enhance paraquat sensitivity of *pry-1(mu38)* animals (**Figure 8F**), which could be explained by significantly reduced expression of *picd-1* in *pry-1* mutants. Finally, as expected from *bh40* being a stronger loss of function allele, the responses of *picd-1(bh40)* animals to chemical exposures were more pronounced than *picd-1(gk3701)*.

Since increased stress sensitivity can affect lifespan and *pry-1* mutants are short lived, we analyzed whether *picd-1* plays a role in aging. The results showed that neither *picd-1*(gk3701) nor *picd-1*(RNAi) enhanced lifespan defects of *pry-1* mutants (**Figure 9A and B, Table 2**). It may be that further knockdown of *picd-1* is unable to exacerbate the phenotype of short-lived *pry-1* mutant animals due to reduced *picd-1* expression as mentioned above. Alternatively, it is plausible that *picd-1* is not involved in lifespan maintenance. To investigate this further, we examined the lifespan of *picd-1* mutant and RNAi-treated animals. The results showed that both *gk3701* and *bh40* alleles caused animals to be short lived. While *picd-1(bh40)* worms showed a significantly reduced lifespan at both 20 °C and 25 °C, such a defect was only seen at 25 °C for *picd-1(gk3701)* animals (**Figures 9B and C, Table 2**). The results are also supported by RNAi experiments. The analysis of age-associated biomarkers revealed a progressive age-associated decline in both the body bending and pharyngeal pumping rates (**Figures 9E and F**). Overall, the data suggest that *picd-1* plays an essential role in maintaining the normal lifespan of animals.

**Table 2:** Lifespan analysis of animals. Each lifespan assay was carried out in two or more batches (see Methods). N: number of animals examined, ns: not significant.

| Genotype | treatment | Mean Lifespan (days) | Median Lifespan (days) | Maximum Lifespan (days) | N | *p* value |
| --- | --- | --- | --- | --- | --- | --- |
| *N2* | - | 16.9 ± 0.9 | 16 | 23 | 56 |  |
| *pry-1(gk3682)* | - | 3.7 ± 0.1 | 4 | 4 | 45 | <0.001 |
| *pry-1(gk3681); picd-1(gk3701)* | - | 3.6 ± 0.1 | 4 | 4 | 50 | <0.001 |
| *picd-1(gk3701)* | - | 15.3 ± 0.5 | 16 | 21 | 44 | ns |
| *picd-1(bh40)* | - | 13.7 ± 0.5 | 14 | 21 | 51 | <0.001 |
| *pry-1(mu38)* | *empty vector* | 3.1 ± 0.3 | 3 | 7 | 79 |  |
|  | *picd-1 RNAi* | 2.9 ± 0.3 | 2 | 8 | 80 | ns |
| *N2* | *empty vector* | 16.6 ± 0.9 | 16 | 21 | 75 |  |
|  | *picd-1 RNAi* | 11.6 ± 0.6 | 18 | 22 | 58 | <0.001 |
| *pop-1(hu9)* | *empty vector* | 11.7 ± 0.8 | 13 | 16 | 54 |  |
|  | *picd-1 RNAi* | 10.1 ± 0.4 | 11 | 13 | 86 | <0.001 |
| *N2* | 25 °C | 13.4 ± 0.5 | 14 | 20 | 56 |  |
| *picd-1(gk3701)* | 25 °C | 11.9 ± 0.5 | 12 | 16 | 48 | <0.001 |
| *picd-1(bh40)* | 25 °C | 10.9 ± 0.4 | 11 | 17 | 54 | <0.001 |

**Figure 9:**
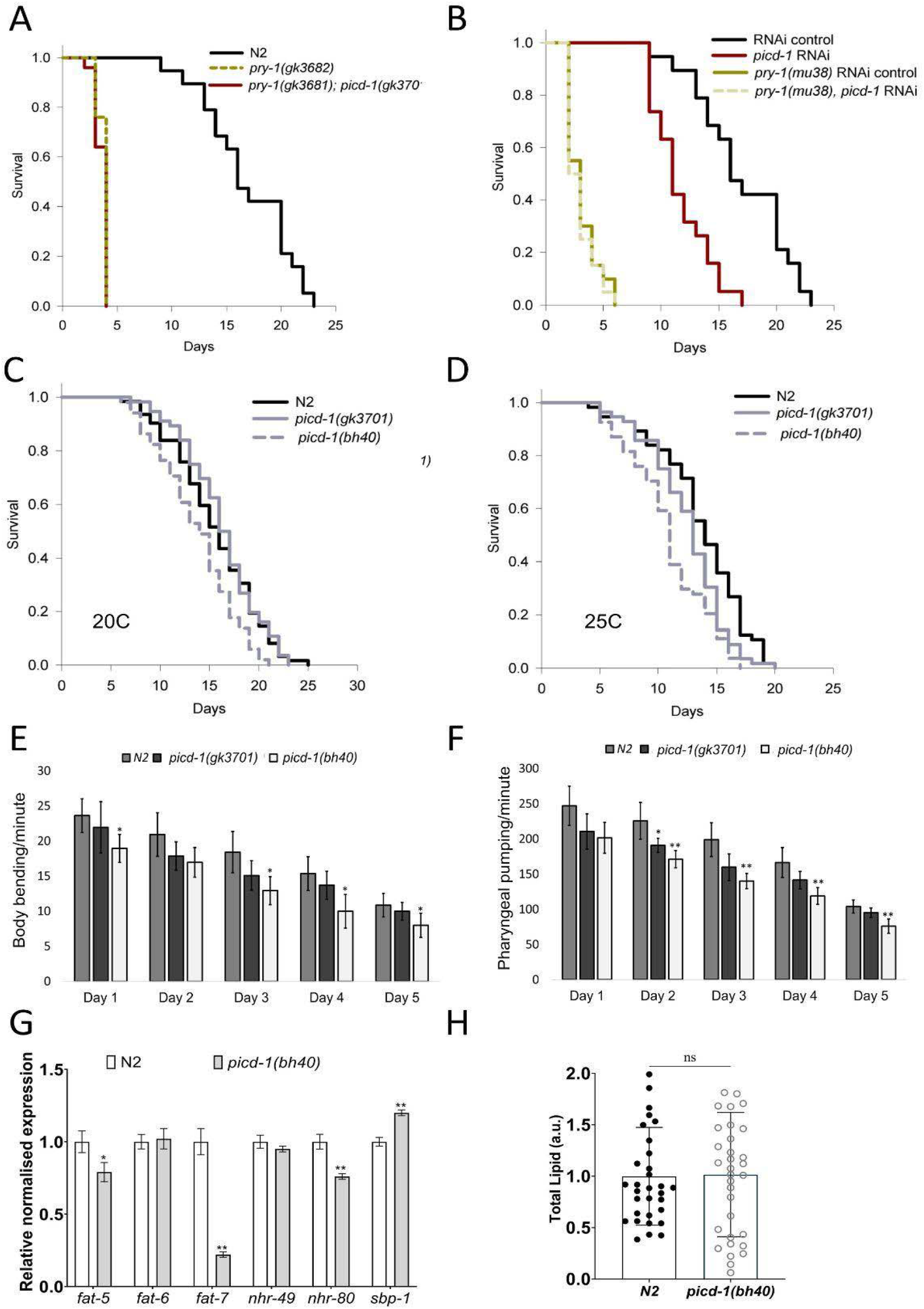
*picd-1* mutation reduces lifespan and shows age-associated deterioration. **(A)** *picd-1* mutation does not affect lifespan of *pry-1* mutants. **(B)** *picd-1* RNAi reduces lifespan of control animals but not that of *pry-1* mutants. **(C-D)** Lifespan of *picd-1(gk3701)* and *picd-1(bh40)* mutants at 20C and 25C. See Materials and Methods section and Table 2 for life span data and statistical analyses. **(E-F)** Bar graphs showing rate of body bending and pharyngeal pumping of *picd-1* mutants compared to wild-type animals over a period of 5 days. Data represent a cumulative of two replicates (n > 15 animals) and error bars represent the standard deviation. Statistical analyses was done using one-way ANOVA with Dunnett’s post hoc test and significant differences are indicated by stars (*): * (*p* <0.05), ** (*p* <0.01). **(G)** Expression analysis of fat-5, fat-6, fat-7, nhr-49, nhr-80 and sbp-1 genes in the *picd-1(bh40)* mutants compared to wild-type. Data represent the mean of two replicates and error bars represent the standard error of mean. Significance was calculated using Bio-Rad software (one-way ANOVA) and significant differences are indicated by stars (*): ** (*p* <0.01). **(H)** Quantification of total lipid using Oil Red O in the wild-type and *picd-1(bh40)* animals. data represent a cumulative of two replicates (n > 30 animals) and error bars represent the standard deviation. Statistical analysis was done using one-way ANOVA with Dunnett’s post hoc test and significant differences are indicated by stars.

In addition to lifespan defects, we showed earlier that PRY-1 regulates lipid metabolism^7^. This prompted us to analyze whether *picd-1* affects lipid levels and expression of genes involved in fatty acid synthesis. The analysis of Δ9 desaturases showed that while *fat-5* and *fat-7* are down, *fat-6* is unaffected (**Figure 9G**). Among the three transcription factors that regulate expression of Δ9 desaturases, *nhr-80* (NHR family) is downregulated, but *sbp-1* (SREBP1 homolog) levels are up (**Figure 9G**)^19^. These results suggest that *picd-1* is needed for normal expression of a subset of lipid synthesis genes. We also quantified lipids by Oil Red O staining and saw no change in *picd-1* mutants (**Figure 9H**), possibly due to functional redundancies within the *fat* family ^20,21^ and *nhr* family of genes^19^. We conclude that *picd-1* is not involved in *pry-1*-mediated lipid regulation.

### Loss of *picd-1* promotes CRTC-1 nuclear localization and upregulates CRTC-1 target genes

Research in *C. elegans* has shown that calcineurin (a calcium-activated phosphatase) signaling promotes nuclear localization of the CREB-regulated transcriptional coactivator (CRTC) homolog CRTC-1, leading to a reduction in lifespan^14^. Given that human CABIN1 negatively regulates calcineurin signaling^10,22^, we investigated whether knocking down *picd-1* could affect the subcellular localization of a translational fusion protein CRTC-1::RFP. The results revealed that RNAi knockdown of *picd-1* caused nuclear localization of CRTC-1, which is consistent with short lifespan of *picd-1* mutants (**Figures 9C-D and 10A**). Moreover, expression of two CRTC-1 responsive genes, *dod-24* and *asp-12* ^23^, was found to be significantly upregulated in *picd-1(bh40)* mutants (**Figure 10B**). Altogether, the data suggest that PICD-1 inhibits CRTC-1 function to regulate the lifespan of *C. elegans.*

**Figure 10:**
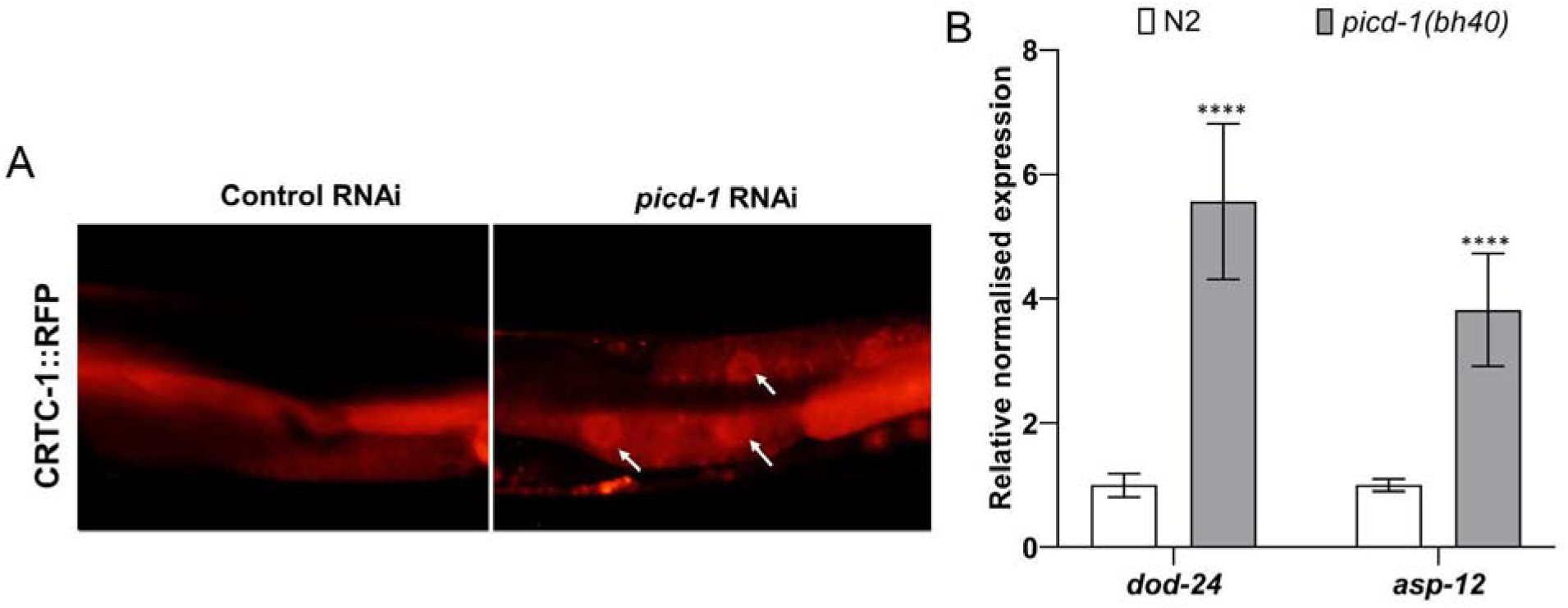
Loss or reduced *picd-1* function promotes CRTC-1 dependent transcriptional response. **(A)** *picd-1* RNAi in the CRTC-1::RFP transgenic animals causes nuclear accumulation of CRTC-1 compared to diffused expression following control (L4440) RNAi. **(B)** qPCR analysis of *dod-24* and *asp-12* in *picd-1(bh40)* animals shows increased expression. Data represent the mean of two replicates and error bars represent the standard error of mean. Significance was calculated using Bio-Rad software (one-way ANOVA) and significant differences are indicated by stars (*): **** (*p* <0.0001).

## DISCUSSION

We have identified a new gene, *picd-1*, in *C. elegans* that interacts with *pry-1* and plays essential roles in multiple larval and adult processes. *picd-1* is predicted to encode a nuclear protein containing a conserved CABIN1 domain. In humans, CABIN1 is a member of the HUCA histone chaperone complex (HIRA/UBN1/CABIN1/ASF1a)^24^. The HUCA complex is implicated in diverse chromatin regulatory events where it preferentially deposits a histone variant H3.3 leading to transcriptional activation by nucleosome destabilization or transcriptional repression through heterochromatinization^25^. Consistent with its important role, CABIN1 is expressed broadly in all human tissues with subcellular localization in the nucleoplasm and cytoplasm^26,27^. Work in other systems has also uncovered homologous proteins. For example, the yeast *S. cerevisiae* contains Hir1p and Hir2p (both HIRA orthologs) and three proteins namely Hir3, Hpc2, and Asf1p that are orthologs of CABIN1, Ubinuclein (UBN1), and ASF1a, respectively^25^.

Our study provides the first genetic evidence of a CABIN1 domain-containing protein in regulating biological processes in *C. elegans*. Other complex components in worms include HIRA-1 (HIRA homolog), ASFL-1, and UNC-85 (both ASF1a homologs)^28–30^. We have shown that mutations in *picd-1* lead to multiple defects such as Pvl, Egl, low brood size, developmental delay, stress sensitivity and short lifespan. Interestingly, loss of *picd-1* function enhances various phenotypes of *pry-1* mutants, some of which are not seen in the *picd-1* mutant alone. For example, *pry-1* and *picd-1* double mutants are Pvl and exhibit P11.p cell fate changes. Also, *picd-1* RNAi enhances seam cell defects in *pry-1* mutant animals. Interestingly, mutations in *picd-1* do not enhance VPC induction and Muv phenotypes of *pry-1* mutants. Overall, these results suggest that *picd-1* participates in a subset of *pry-1*-mediated processes.

We also analyzed the role of *picd-1* in other PRY-1-mediated non-developmental events such as egg-laying, embryonic survivability, aging, stress response and lipid metabolism. Loss of *picd-1* function worsened the embryonic lethality of *pry-1* mutants. Moreover, *pry-1; picd-1* double mutants have very low brood size due to defects in gonad arms. Similar phenotypes are seen in mutants of other HIRA complex components. Thus, knockdown of *hira-1* leads to embryonic lethality, *asfl-1* or *unc-85* single mutants have low brood size, and *asfl-1; unc-85* double mutants are sterile^29–31^. Together, these data show that *pry-1* and *picd-1* interact to regulate embryonic viability and fertility of animals. However, it remains to be seen whether PRY-1 and PICD-1 interact with other HIRA complex components to mediate their function.

Among other roles, we found that *picd-1* is needed for normal stress response maintenance. Specifically, *picd-1* mutants show enhanced sensitivity to paraquat and tunicamycin. Consistent with this, mutant animals exhibit increased levels of UPR markers. Both the *picd-1* and *pry-1* mutants significantly increase *hsp-16.2,* and *hsp-4*, suggesting that the genes are involved in regulating ER-UPR and HSR. More work is needed to determine whether the two genes uniquely affect MT-UPR and oxidative stress, and their biological significance.

Mutants that show sensitivity to stress are often short lived^32–34^. The analysis of lifespan phenotype revealed that similar to *pry-1* mutants, *picd-1(bh40)* animals are short lived and exhibit defects in age-related physiological markers, which is consistent with both genes functioning together to regulate stress response and aging. However, there are functional differences between them. For example, lipid levels are greatly reduced in *pry-1* mutants but unaffected in *picd-1* mutants. We found that *nhr-80* and *fat-7* levels are reduced in *picd-1* mutant animals, which is consistent with the known role of *nhr-80* regulating *fat-7* expression^21^. Overall, while *picd-1* is needed for normal expression of *fat-5*, *fat-7* and *nhr-80*, a lack of its function does not compromise lipid levels in animals. The differences between *pry-1* and *picd-1* with respect to lipid regulation suggest that *picd-1* participates only in a subset of *pry-1*-mediated processes. However, to what extent the two genes interact in specific tissues and the precise nature of their interactions is unknown.

A possible mechanism of *picd-1* function in lifespan maintenance may utilize calcineurin. AMPK and calcineurin modulation of CRTCs is conserved in mammals and *C. elegans*^14^. In *C. elegans*, AAK-2 and calcineurin regulate CRTC-1 post-translationally in an opposite manner, where activated AAK-2 causes nuclear exclusion of CRTC-1 and extends lifespan. Such a phenotype is also seen after deactivating calcineurin^14^. Our data shows that loss of *picd-1* function causes nuclear localization of CRTC-1 and activates the expression of *crtc-1* target genes. These findings together with the fact that mammalian CABIN1 inhibits calcineurin-mediated signaling^10,22,35^, suggests that PICD-1 may regulate CRTC-1 via downregulation of calcineurin in *C. elegans.* As the loss of *picd-1*/CABIN-1 should lead to increased calcineurin signaling, this may explain the shorter lifespan of *picd-1* mutants. Given that *picd-1* is downregulated in *pry-1* mutants, and both genes are needed to delay aging and confer stress resistance, it suggests that *pry-1* and *picd-1* may interact to maintain stress response and lifespan of animals. There are many unanswered questions, for example, whether *picd-1* is regulated by *pry-1* in a Wnt dependent manner during aging and if such a mechanism involves *pop-1*, as well as if both stress response and lifespan processes are regulated by a common set of genes acting downstream of *pry-1* and *picd-1*. Additionally, it is unclear if other HUCA complex components regulate lifespan and stress response by utilizing PICD-1 and PRY-1. Further work is needed to investigate these questions and to gain a deeper understanding of conserved mechanisms involving AXIN and CABIN1 function in eukaryotes.

## MATERIALS AND METHODS

### Worm strains

Animals were maintained at 20 °C on standard nematode growth media (NGM) plates seeded with OP50 *E. coli* bacteria.

N2 (wild-type *C. elegans*)

DY220 *pry-1(mu38)*

VC3710 *pry-1(gk3682)*

VC3709 *pry-1(gk3681); picd-1(gk3701)*

DY725 *pry-1(mu38); picd-1(bh40)*

DY678 *bhEx287[pGLC150(picd-1p::gfp) + myo-3::wCherry]*

DY698 *picd-1(bh40)*

DY694 *picd-1(gk3701)*

RG733 *wIs78[(scm::GFP) + (ajm-1p::GFP)]*

AGD418 *uthIs205[crtc-1p::CRTC-1::RFP::unc-54 3’ UTR + rol-6(su1006)]*

### Mutant allele generation

The *gk3701* allele that was created during the process of creating a CRISPR mutant of *pry-1*, removes 5bp (GGTGA) (flanking 25 nucleotides: GTGAAGAGGATGAGGACAATGGTGA and GGATTCAGAAGAAGAAGATGAAGAA) of the second exon. This frameshift mutation leads to a premature stop codon in the second exon followed by multiple consecutive stop codons (See primers used in **Table S2**).

The allele *bh40* was created using the Nested CRISPR technique^36^. Here we replaced 84bp of the first exon with a sequence containing multiple stop codons both in frame and out of frame of the coding transcript (See primers used in **Table S2**). Both mutants have been out crossed twice with the wildtype N2 animals and sequence confirmed.

### RNAi

RNAi mediated gene silencing was performed using a protocol previously published by our laboratory^37^. Plates were seeded with *Escherichia coli* HT115 expressing either dsRNA specific to candidate genes or empty vector (L4440). Synchronized gravid adults were bleached, and eggs were plated. After becoming young adult animals were analyzed for vulva or seam cell phenotype.

### Fluorescent microscopy

Animals were paralyzed in 10mM Sodium Azide and mounted on glass slides with 2% agar pads and covered with glass coverslips for immediate image acquisition using Zeiss Apotome microscope and software.

### Vulval induction, P12.pa, body bending, and pharyngeal pumping

Vulva phenotype in L4 stage animals was scored using a Nomarski microscope. VPCs were considered as induced if they contained progeny. Three VPCs are induced in Wild-type (N2) animals. Mutants carrying more than 3 induced VPCs were termed as ‘over-induced’. Muv and Pvl phenotypes were scored in adults.

P12.pa cell fates were quantified in L4 stage animals. Wild-type (N2) larvae have one P11.p and P12.pa cells. In *pry-1* mutants, two P12.pa-like cells are observed and P11.p is missing.

Rate of body bending per 1 min and the rate of pharyngeal pumping per 30 sec for adults were analyzed over the period of 4 days^6^. Hermaphrodites were analyzed for these phenotypes under the dissecting microscope in isolation on OP50 plates. Pharyngeal pumping was assessed by observing the number of pharyngeal contractions for 30 sec. For body bending assessment, animals were stimulated by tapping once on the tail of the worm using the platinum wire pick where one body bend corresponded to one complete sinusoidal wave of the worm. Only animals that moved throughout the duration of 1 min were included in the analysis.

### Lifespan analysis

Lifespan experiments were done following adult specific RNAi treatment using a previously described protocol^8^. Animals were grown on NGM OP50 seeded plates till late L4 stage after which they were transferred to RNAi plates. Plates were then screened daily for dead animals and surviving worms were transferred every other day till the progeny production ceased. Censoring was done for animals that either escaped, burrowed into the medium, showed a bursting of intestine from the vulva or underwent bagging of worms (larvae hatches inside the worm and the mother dies)^38^.

### Stress assay

Oxidative (paraquat) and endoplasmic reticulum mediated stress (tunicamycin) stress experiments were performed using 100mM paraquat (PQ) (Thermo Fisher Scientific, USA) and 25ng/μL tunicamycin (Sigma-Aldrich, Canada) respectively. Animals were incubated for 1hr, 2hr, 3hr and 4hr, following the previous published protocol^6^. All the final working concentrations were made in M9 instead of water. At least 30 animals were tested for each strain in each replicate. Mean and standard deviation were determined from experiments performed in duplicate. Animals were considered dead if they had no response following a touch using the platinum wire pick and showed no thrashing or swimming movement in M9. Moreover, dead animals usually had an uncurled and straight body shape in comparison to the normal sinusoidal shape of worms.

### Oil Red O staining

Neutral lipid staining was done on synchronized day-1 adult animals using Oil Red O dye (Thermo Fisher Scientific, USA) following the previously published protocol^9^. Quantification was then done using ImageJ software as described previously^39^.

### Molecular Biology

RNA was extracted from synchronized L3 and day-1 adult animals. Protocols for RNA extraction, cDNA synthesis and qPCR were described earlier^7^. Briefly, total RNA was extracted using Trizol (Thermo Fisher, USA), cDNA was synthesized using the SensiFast cDNA synthesis kit (Bioline, USA), and qPCR was done using the SYBR green mix (Bio-Rad, Canada). Primers used for qPCR experiments are listed in **Table S1**.

### Statistical analyses

Statistics analyses were performed using GraphPad prism 9, SigmaPlot software 11, CFX Maestro 3.1 and Microsoft Office Excel 2019. For lifespan data, survival curves were estimated using the Kaplan- Meier test and differences among groups were assessed using the log-rank test. qPCR data was analyzed using Bio-Rad CFX Maestro 3.1 software. For all other assays, data from repeat experiments were pooled and analyzed together and statistical analyses were done using GraphPad Prism 9. *p* values less than 0.05 were considered statistically significant.

## ACKNOWLEDGEMENT

We thank Hannan Minhas for his help with the electrotaxis experiments and Wouter van den Berg for his help during the screening of CRISPR mutant allele *picd-1(bh40)*. This work was supported by NSERC Discovery grant to BG and NSERC CGS-D scholarship to AM. Some of the strains were obtained by the *Caenorhabditis* Genetics Center (CGC), which is funded by NIH Office of Research Infrastructure Programs (P40 OD010440).

## AUTHORS CONTRIBUTIONS

AM initially characterized *picd-1* mutants and generated many reagents for the study. AM, SM and SKBT performed several experiments. AM, SKBT, and BG analyzed data. BG supervised the study.

